# A DOT1B/Ribonuclease H2 protein complex is involved in R-loop processing, genomic integrity and antigenic variation in *Trypanosoma brucei*

**DOI:** 10.1101/2020.03.02.969337

**Authors:** Nicole Eisenhuth, Tim Vellmer, Falk Butter, Christian J. Janzen

**Author notes:** To whom correspondence should be addressed. Christian J. Janzen, Falk Butter.

## Abstract

The parasite *Trypanosoma brucei* periodically changes the expression of protective variant surface glycoproteins (VSGs) to evade its host’s immune system in a process known as antigenic variation. One route to change VSG expression is the transcriptional activation of a previously silent VSG expression site (ES), a subtelomeric region containing the *VSG* genes. Homologous recombination of a different *VSG* from a large reservoir into the active ES represents another route. The conserved histone methyltransferase DOT1B is involved in transcriptional silencing of inactive ES and influences ES switching kinetics. The molecular machinery that enables DOT1B to execute these regulatory functions remains elusive, however. To better understand DOT1B-mediated regulatory processes, we purified DOT1B-associated proteins using complementary biochemical approaches. We identified several novel DOT1B-interactors. One of these was the Ribonuclease H2 complex, previously shown to resolve RNA-DNA hybrids, maintain genome integrity, and play a role in antigenic variation. Our study revealed that DOT1B depletion results in an increase in RNA-DNA hybrids, accumulation of DNA damage and recombination-based ES switching events. Surprisingly, a similar pattern of VSG deregulation was observed in Ribonuclease H2 mutants. We propose that both proteins act together in resolving R-loops to ensure genome integrity and contribute to the tightly-regulated process of antigenic variation.

## INTRODUCTION

Antigenic variation is one of the most sophisticated strategies used by protist parasites, such as *Trypanosoma brucei*, to escape the immune system of their mammalian hosts (1). A prerequisite for this process is the periodic exchange of the trypanosomes’ surface coat, composed of densely packed proteins called variant surface glycoproteins (VSGs). Even though trypanosomes possess a large repertoire with more than 2,000 *VSG* genes and pseudogenes located in subtelomeric regions of their eleven megabase- and approximately five intermediate- and 100 minichromosomes (2,3), only one *VSG* is expressed at any given time from one out of 15 subtelomeric expression sites (ES) (4). Switching of the VSG coat can be facilitated either by transcriptional activation of a previously-silent ES, a so-called *in situ* switch, or by recombination of another *VSG* gene into the active ES.

DNA recombination is the major route of VSG switching and ensures that the full repertoire of *VSGs* from subtelomeric genome loci can be used (5). DNA accessibility and genome architecture influenced by histone variants H3.V and H4.V impact recombination events in the ES (6). Furthermore, the interplay of several proteins important for homologous recombination are required for this process (7), and it has become clear that DNA lesions trigger recombination events (8,9). Different possibilities for a source of recombination-initiating DNA breaks have been proposed, and a recently published model has suggested the participation of RNA-DNA hybrids (so-called R-loops) (10). R-loops in *T. brucei* are generated by transcription (11), and usually removed by the two types of Ribonuclease H (RH1 and RH2) (12-14). DNA lesions and altered *VSG* expression were observed if R-loops were not processed properly (12,13), or if their formation at the ES was not accurately coordinated by the telomere-associated protein RAP1 (15).

The monoallelic expression of *VSG* from one ES and the coordinated switching to another ES is tightly controlled at multiple levels (16). Chromatin and chromatin-associated factors play ubiquitous roles in *VSG* expression control (17). The chromatin structure of an active ES is distinct from a silent one: the active ES is depleted of nucleosomes (18,19), and is instead enriched for the high mobility group box protein TDP1 (20). The active ES is transcribed from an unusual extranucleolar locus called the Expression Site Body (ESB) by RNA polymerase I (RNA pol I) (21). Proteins associated with this ESB such as the class I transcription factor A (CITFA) complex or the VSG-exclusion (VEX) complex are required for RNA pol I recruitment and monoallelic transcription, respectively (22,23). Interestingly, R-loops also seem to have an impact on transcriptional control; after depletion of monomeric RH1 or the catalytic subunit of trimeric RH2, trypanosomes derepress silent ESs and express different VSGs on their surface (12,13). Furthermore, transcriptional control is also regulated by nucleosome assembly and chromatin remodeling at the promoter region of the ES, mediated by the ISWI (24), FACT (25), CAF-1b, and ASF1A protein complexes (26). Not only the positioning of nucleosomes, but also their post-translational modifications (PTMs) play a role in ES regulation. For example, the conserved histone methyltransferase DOT1B impacts ES expression on multiple levels (27).

DOT1 and its mono-, di-, and trimethylation of histone H3 lysine 79 (H3K79me1/2/3) functions in several nuclear processes in eukaryotes, including transcription, the DNA damage response, and telomeric silencing (28). Knowledge of DOT1 interactions with other H3K79 readers has provided valuable insights into these functions. For instance, competition between H3K79 methylation by yeast Dot1p and the binding of silent information regulator (Sir) proteins to the same chromatin target are involved in the formation of heterochromatin at telomeres (29,30). Additionally, H3K79 methylation is associated with actively-transcribed genes (31). The interaction between human DOT1L and the phosphorylated C-terminal domain of RNA pol II could provide a recruiting mechanism of DOT1L to such active genes (32). DOT1L also interacts directly with members of the AF10 and ENL protein families (33-35), which are usually found in RNA pol II transcriptional elongation complexes (36-39). Incorrect DOT1L recruitment as a result of chimeric gene fusions between DOT1L-interacting proteins AF10/ENL and the histone H4 methyltransferase Mixed Lineage Leukemia (MLL) lead to aberrant H3K79 methylation patterns and increased transcription of oncogenes that cause leukemia (35,40,41), further supporting a role of DOT1B in transcription. DOT1 also helps to maintain genome integrity in mitotic cells by acting through a variety of repair pathways after DNA damage has occurred (28). For instance, binding of the checkpoint adaptor Rad9 to methylated H3K79 in yeast regulates the resectioning step necessary for repair of DNA double strand breaks (DSBs) by homologous recombination (42). The binding of the Rad9 homolog 53BP1 to methylated H3K79 is also important for detection and repair of DSBs in humans (43), and is further promoted through an interaction of DOT1L to the HLA-B-associated transcript Bat3 (44). Additionally, Dot1p inhibits the error-prone polymerases of the translesion synthesis pathway in response to DNA damage caused by alkylating agents (45). Dot1p is further required for the response to meiotic DSBs and modulates the meiotic checkpoint response (46,47). DOT1 requires a nucleosomal context for its catalytic activity (48) with multivalent interactions to DNA, H2A, H2B, H3 (49,50) and the H4 tail (51). The methylation activity of DOT1 is stimulated by ubiquitination of H2B (52,53). This ubiquitin mark reduces the sampling space of DOT1L on the nucleosome by conformational restriction, which enables DOT1L to catalyze higher methylation states (50). Further work is needed to better understand the exact mechanisms and relative contributions of different regulators and interactors of DOT1.

In contrast to yeast and mammals, trypanosomes possess two DOT1 paralogs, DOT1A and DOT1B, with different enzymatic activities. The mono- and dimethylation of histone H3 on lysine 76 (H3K76) is mediated by DOT1A and is essential for replication initiation (54). In contrast, trimethylation of H3K76 is catalyzed by DOT1B (55). DOT1B knockout (KO) causes a defect in the differentiation from the mammalian-infective stage to the insect-infective stage (56), which is accompanied by severe karyokinesis defects (57). In addition, DOT1B plays an important role in the regulation of *VSG* transcription: KO cells show derepression of transcriptionally silent ES *VSG*s and extremely slow ES *in situ* switch kinetics (27). Furthermore, the attenuation of the active ES in response to inducible expression of an additional *VSG* gene is not observed in DOT1B-negative parasites (58). The molecular machinery which enables DOT1B to execute regulatory functions at the ES is still elusive, however.

To fully understand how DOT1B can influence these different nuclear processes in *T. brucei*, we purified DOT1B protein complexes using complementary biochemical approaches. Surprisingly, one of the most abundant DOT1B-associated protein complexes was RH2, which is important for the maintenance of genome integrity by resolving R-loops (13). Since a contribution of RH2 to antigenic variation was shown recently (13), we predicted that DOT1B could support this function. Consistent with this hypothesis, we found that DOT1B depletion caused an increased R-loop abundance, accumulation of DNA damage, and aberrant *VSG* expression throughout the *VSG* repertoire, which indicates additional recombination events at the ESs.

## MATERIAL AND METHODS

### Trypanosome cell lines and cultivation

*T. brucei* BSFs (Lister strain 427, antigenic type MITat 1.2, clone 221a) were maintained in HMI-9 medium with 10% heat-inactivated fetal calf serum (Sigma) at 37°C and 5% CO_2_ (59). PCF trypanosomes (Lister strain 427) were cultured in modified SDM-79 medium with 10% heat-inactivated fetal calf serum (Sigma) at 27°C and 5% CO_2_ (60). Transgenic BSF “2T1” (61), BSF “Single Marker” (SM) and PCF “29-13” derivatives which constitutively express a T7 RNA polymerase and a Tet repressor (62) were used for regulated expression experiments. Cell densities were determined using a Coulter Counter Z2 particle counter (Beckman Coulter) and cultures were maintained in exponential growth phase by regular dilution. Transfections and drug selections of trypanosomes were carried out as described previously (63).

#### PTP::DOT1B / Δdot1b

The first allele of *DOT1B* (Tb427.1.570) was tagged endogenously at the 5’ end with *PTP* as previously published (64). Briefly, a fragment of the *DOT1B* ORF (position 1-783) was amplified from genomic DNA and cloned into the p2678_PUR_PTP plasmid. NsiI was used to linearize the vector within the *DOT1B* fragment prior to transfection of BSF and PCF cells. A PCR-based gene deletion approach was used to replace the second *DOT1B* allele with the hygromycin phophotranspherase (*HYG*) ORF in BSFs or with the blasticidin-S-deaminase (*BSD*) ORF in PCFs. *HYG* ORF was amplified from the pHD309 plasmid and *BSD* ORF was amplified from the pPOTv4 plasmid with primers containing 60 bp homology-flanks to the *DOT1B* UTRs for recombination. Correct integration of the constructs was verified by PCR using primers annealing in the UTRs of *DOT1B* and ORFs of the resistance cassettes.

#### PTP^Ti^

To express the *PTP* tag ectopically, its ORF was amplified from p2678 with forward primers containing a HindIII restriction site and reverse primers containing a stop codon and BamHI site. The *PTP* internal BamHI site was destroyed by site-directed mutagenesis and the construct was cloned into the pLEW100v5bd plasmid. The plasmid was linearized with NotI prior to transfection of 29-13 PCF cells. Expression was induced with 50 ng/ml tetracycline for 24h.

#### RH2A::HA, RH2B::HA and RH2C::HA

Both alleles of *RH2A* (Tb427.10.5070), *RH2B* (Tb427.1.4220) and *RH2C* (Tb427.1.4730) were in situ-tagged with a 3xHA epitope sequence at the 3’ end using a PCR-based method and the pMOTag2H and pMOTag4H plasmids as previously described (65). Correct integrations of constructs was tested by using primers annealing in the UTRs of the genes of interest and the ORFs of resistance cassettes.

#### DOT1B-Myc-BirA^Ti^

For inducible expression of the DOT1B-Myc-BirA* fusion protein, the *DOT1B* ORF (without stop codon) was amplified from genomic DNA and cloned into the pLEW100_Myc_BirA* plasmid (66) between HindIII and NdeI sites. The plasmid was linearized with NotI prior to transfection of 29-13 PCF cells. Expression was induced with 5 ng/ml tetracycline.

#### DOT1B and RH2A RNAi

RNAi targets (*RH2A*: position 14-398 of 981 bp ORF; *DOT1B*: position 132-480 of 828 bp ORF) were amplified from genomic DNA with primers containing attB1 sites for BP recombination (Invitrogen) into the stem-loop RNAi vector pGL2084 (67). 2T1 cells were transfected with SgsI-linearized plasmids and RNAi of clones with correctly integrated constructs was induced with 1 µg/ml tetracycline.

### IgG affinity purification

IgG affinity purification was carried out as previously described (68), with minor changes: four biological replicates per cell line (1×10^8^ PCF cells and 4×10^8^ BSF cells per replicate) were harvested by centrifugation (1,500 × g, 10 min, 4°C), washed in 10 ml ice-cold wash solution [20 mM Tris-HCl pH 7.4, 100 mM NaCl, 3 mM MgCl_2_, 1 mM EDTA] and 10 ml ice-cold extraction buffer [150 mM sucrose, 150 mM KCl, 20 mM potassium L-glutamate, 3 mM MgCl_2_, 20 mM HEPES-KOH pH 7.7, 0.1% Tween 20, 1 mM dithiothreitol (DTT), 10 μg/ml leupeptin, 10 μg/ml aprotinin, EDTA-free protease inhibitor cocktail (Roche)]. Cells were lysed in 1 ml extraction buffer by three freeze-thaw cycles using liquid nitrogen and by sonication using a Bioruptor (Diagenode) with one 30 s high power pulse. The solubilized material was cleared by centrifugation (20,000 × g, 10 min, 4°C) and supernatants were stored at 4°C. Per TAP, 20 µl IgG Sepharose 6 Fast Flow beads (GE Healthcare) were washed twice with 1 ml ice-cold PA-150 buffer [150 mM KCl, 20 mM Tris-HCl pH 7.7, 3 mM MgCl_2_, 0.5 mM DTT, 0.1% Tween20] (500 × g, 5 min, 4°C) and added to the cleared lysates. The lysate beads mixture was incubated with rotary mixing for 2 h at 4°C. Beads were washed twice with 1 ml PA-150 buffer containing EDTA-free protease inhibitor cocktail (Roche) (500 × g, 5 min, 4°C) and proteins were eluted by heating in 65 µl NuPAGE LDS sample buffer (Thermo Fisher) supplemented with 100 mM DTT (10 min, 70°C). Beads were pelleted (1,000 × g, 1 min, RT) and the supernatant was transferred with a Hamilton syringe to a new tube.

### Co-Immunoprecipitation

30 µl of Protein G Sepharose Fast Flow bead slurry (GE Healthcare) was washed once with 1 ml ice-cold phosphate-buffered saline (PBS) (500 × g, 1 min, 4°C) and twice with 1 ml ice-cold 1% bovine serum albumin (BSA)/PBS. Beads were blocked by rotating in 1 ml 1% BSA/PBS (1 h at 4°C). After harvesting the beads by centrifugation (500 × g, 1 min, 4°C), 50 µl anti-HA/12CA5 monoclonal mouse IgG antibody was bound to the beads in a final volume of 1 ml PBS (overnight, 4°C, rotary mixing). Three washing steps with 1 ml ice-cold 0.1% BSA/PBS removed unbound antibody. The antibody-sepharose conjugate was stored at 4°C. Quadruplets of 1×10^9^ cells per cell line were harvested (1,500 × g, 10 min, 4 °C) and washed in 10 ml ice-cold Trypanosome Dilution Buffer (TDB) [5 mM KCl, 80 mM NaCl, 1 mM MgSO_4_, 20 mM Na_2_PO_4_, 2 mM NaH_2_PO_4_, 20 mM glucose, pH 7.4]. Cells were lysed by incubation in 1 ml ice-cold IP buffer [150 mM NaCl, 20 mM Tris-HCl pH 8.0, 10 mM MgCl_2_, 0.25% Igepal CA-630, 1 mM DTT, EDTA-free Protease Inhibitor Cocktail (Roche)] for 20 min on ice and by sonication (3 cycles, 30 sec on, 30 sec off, settings high). Lysates were cleared by centrifugation (20,000 × g, 10 min, 4 °C). Soluble proteins were incubated with 30 μl antibody-sepharose conjugate, which was previously equilibrated in 1 ml IP buffer (500 × g, 5 min, 4°C). After an incubation step under mild agitation (3h, 4°C), beads were washed twice with 1 ml IP buffer (5 min, on ice) and harvested by centrifugation (500 × g, 5 min, 4°C). Bound proteins were eluted as described above.

### BioID

BioID was conducted as described in (66) with minor changes. Expression of DOT1B-Myc-BirA* was induced by addition of 5 ng/ml tetracycline for 24 h and cells were further incubated for 24 h with 50 µM D-biotin (Invitrogen) and 5 ng/ml tetracycline. Quadruplets of 1×10^9^ DOT1B-Myc-BirA*-expressing cells and of the uninduced control cells were harvested by centrifugation (1,800 × g, 5 min, 4°C). After washing cells three times with PBS, the cells were incubated in 1 ml IP buffer (20,min, on ice) and then sonicated (6 cycles, 30 sec on, 30 sec off, settings high). Lysates were cleared by centrifugation (15,000 × g, 10 min, 4°C) and further stored on ice. Pierce Streptavidin Agarose beads (Thermo Fisher) were washed twice with 1 ml ice-cold binding buffer [50 mM Na_2_HPO_4_, 50 mM NaH_2_PO_4_, 150 mM NaCl] and equilibrated with 1 ml ice-cold IP buffer (500 × g, 1 min, 4°C). 50 µl beads were added to each cleared lysate and the reactions were incubated under mild agitation (4h, 4°C). The unbound material was separated from the beads by centrifugation (500 × g, 5 min, 4°C) and was followed by two washing steps with 1 ml IP buffer (5 min, on ice). Bound proteins were eluted as described above.

### Mass spectrometry and data analysis

Samples were run on a Novex Bis-Tris 4-12% gradient gel (Thermo Fisher) with MOPS buffer (Thermo Fisher) for 10 min at 180V. The gel was stained with coomassie blue G250 dye (Biozym). Each lane was cut into pieces, minced and destained in water/50% EtOH. The gel pieces were dehydrated with pure ACN, reduced with 10 mM DTT (Sigma Aldrich) and subsequently alkylated with 55 mM iodoacetamide in the dark. The again dried gel pieces were digested with 1 µmol trypsin at 37°C overnight. The digested peptides were desalted and stored on StageTips (69) until measurement.

Peptides were separated along a 240 min gradient (90 min for VSG analysis) on a EasyLC 1000 uHPLC system using a C18 reverse phase column, packed in-house with Reprosil C18 (Dr. Maisch GmbH). The column was enclosed into a column oven (Sonation) and the peptides were sprayed into a Q Exactive Plus mass spectrometer (Thermo Fisher). The mass spectrometer was operated in a data-dependent acquisition mode using a top10 method. Spray voltage was set at ca. 2.4 kV.

The aquired raw files were analyzed with MaxQuant (version 1.5.8.2) (70) using the *Trypanosoma brucei* protein database downloaded from TriTrypDB. Prior to bioinformatics analysis, contaminants, reverse hits, proteingroups only identified by site and proteingroups with less than 2 peptides (one of them unique) were removed.

### Western blot analysis

Western blots were carried out according to standard protocols. In brief, lysates (including phosphatase inhibitors (Merck) in the case of the quantitative γH2A-assay) of 2×10^6^ cells (lysates of 4×10^6^ cells in the case of RH2::HA cell lines) were separated by SDS-PAGE on 12% - 15% polyacrylamide gels and transferred to PVDF membranes. After blocking membranes (1 h, RT) in indicated solutions, primary antibodies diluted in 0.1% Tween/PBS were incubated with the membrane (1 h, RT). After three washing steps with 10 ml 0.2% Tween/PBS, the membranes were incubated with IRDye 800CW- and 680LT-coupled secondary antibodies or IRDye 680RD Streptavidin (LI-COR Bioscience) in 0.1% Tween/PBS supplemented with 0.02% SDS (1 h, RT). Secondary antibodies were diluted according to the manufacturer’s instructions. Signals were imaged using a LI-COR Odyssey CLx and quantified with ImageStudio software. The following primary antibodies were used in this study: polyclonal rabbit anti-H3K76me2 (1:2,000, blocked in 3% BSA/PBS) (56); polyclonal rabbit anti-H3K76me3 (1:2,000, blocked in 3% BSA/PBS) (56); monoclonal rat anti-DOT1B/13’E5 was a gift from E. Kremmer, Helmholz Centre Munich (1:2,000, blocked in 5% milk/PBS); rabbit anti-γH2A was a gift from R. McCulloch, University of Glasgow (1:2,000, blocked in 3% BSA/PBS); monoclonal mouse anti-HA/12CA5 (1:500, blocked in 5% milk/PBS); polyclonal rabbit anti-TbH3 (1:50,000 dilution, blocked in 5% milk/PBS) (54); monoclonal mouse anti-PFR1,2 (L13D6) is specific for two paraflagellar rod proteins, was used for signal intensity normalization, and was a gift from K. Gull, University of Oxford (1:200 dilution).

### R-loop dot blot

Genomic DNA was isolated from 2.5×10^7^ cells using the High Pure PCR Template Preparation kit (Roche). 1.1 µg DNA was then either treated with 10 U recombinant *E. coli* Ribonuclease H (NEB) or ddH_2_O in 1xRibonuclease H buffer (2 h, 37°C). All samples were further incubated with 10 µg RNase A (Thermo Fisher) (1 h, 37°C) in a final concentration of 500 mM NaCl. The DNA samples were spotted on a Hybond N+ membrane (GE Healthcare) using a dot blot apparatus in a two-fold serial reduction and cross-linked with UV (0.12 J). To quantify R-loop formation, the membrane was blocked in 5% milk/PBS (overnight, 4°C). 1 µg/ml S9.6 antibody (Merck Millipore) in 0.1% Tween/PBS was then incubated with the membrane (1 h, RT). After three washes with 50 ml 0.2% Tween/PBS, the membrane was incubated with HRP-conjugated goat anti-mouse antibody (Dianova) (1:20,000) (1 h, RT). After three washes with 50 ml 0.2% Tween/PBS, the HRP signal was developed with the Western lightning Plus-ECL solutions (PerkinElmer) and imaged using an iBright Imaging System (Thermo Fisher). iBright Analysis software was used for quantification of signals.

### VSG-2 analysis by FACS

Abundance of VSG-2 on the cell surface was analyzed as previously described (27), with minor changes. 5×10^5^ cells were harvested with a pre-cooled centrifuge (1,500 × g, 5 min, 4°C) and resuspended in 150 µl ice-cold HMI-9 medium containing AlexaFluor647-conjugated monoclonal mouse VSG-2 antibody (71) (1:500). After 50 min of incubation at 4°C with gentle rotary mixing at 6 rpm, fluorescence signal of cells was immediately quantified with a FACSCalibur cell analyzer (BD).

### Isolation of soluble VSGs for MS analysis

Soluble VSGs were isolated in quadruplicates from 4×10^7^ cells according to previously published protocols (72). Briefly, cells were pre-cooled (10 min, on ice) prior to harvesting by centrifugation (1,500 × g, 10 min, 4°C). After a washing step with 10 ml ice-cold TDB (1,500 × g, 10 min, 4°C), cells were transferred into a microcentrifuge tube in 1 ml ice-cold TDB and pelleted again (1,500 × g, 10 min, 4°C). Subsequently the pellet was resuspended in 45 µl 10 mM sodium phosphate buffer supplemented with EDTA-free Protease Inhibitor Cocktail (Roche) and incubated (5 min, 37°C). After cooling down cells (2 min, on ice), they were pelleted by centrifugation (14,000 × g, 5 min, 4°C) and the supernatant containing soluble VSGs was transferred into a new reaction tube. After addition of 24.5 µl NuPAGE LDS sample buffer (Thermo Fisher) supplemented with 100 mM DTT, samples were boiled (10 min, 70°C).

## RESULTS

### Identification of novel DOT1B-interacting proteins in trypanosomes

To understand the different DOT1B-dependent processes in trypanosomes, it is essential to identify the molecular components that are involved in these distinct functions. Attempts to purify DOT1B under endogenous expression levels using conventional epitope-based methods were consistently unsuccessful (our unpublished data). We therefore employed a highly-efficient Tandem Affinity Purification (TAP) approach with the improved PTP tag in order to find DOT1B-interacting proteins (68). The *PTP* tag was fused at the 5’ end of one allele of *DOT1B* in the mammalian-infective bloodstream form (BSF) stage of the parasite. The second allele of *DOT1B* was knocked out using homologous recombination (Supplementary Figure S1A). Correct integration of the targeting and KO constructs was confirmed by PCR analysis of genomic DNA (Supplementary Figure S1B). The KO did not significantly affect the enzymatic activity of DOT1B, because the trimethylation level of H3K76 was comparable to that of wild type (WT) cells (Supplementary Figure S1C). Affinity purification was performed with protein extracts of parasites that expressed the PTP-DOT1B fusion protein. Extracts of WT cells served as a control. The purified proteins, along with fractions obtained during the procedure, were analyzed by Western blot (WB) using anti-DOT1B antibodies. The WBs confirmed the successful enrichment of DOT1B (Supplementary Figure S2A).

To elucidate the composition of a potential DOT1B complex, we compared the purified samples of PTP::DOT1B cells with those from WT cells using label-free quantitative proteomics. As expected, DOT1B was significantly enriched (p< 0.05) in the eluate fraction of the PTP-DOT1B BSF pulldown, as well as 22 other proteins (Figure 1A). Three proteins were significantly enriched in the control lysates. The whole dataset is summarized in Supplementary Table S1. As DOT1B has a nuclear localization (73), and acts as a histone methyltransferase on chromatin (56), we further focused on proteins predicted to be nuclear. Four proteins, besides DOT1B were categorized as such based on nuclear proteome data (74), TrypTag (73), and relevant literature (as indicated in Supplementary Table S1).

**Figure 1.**
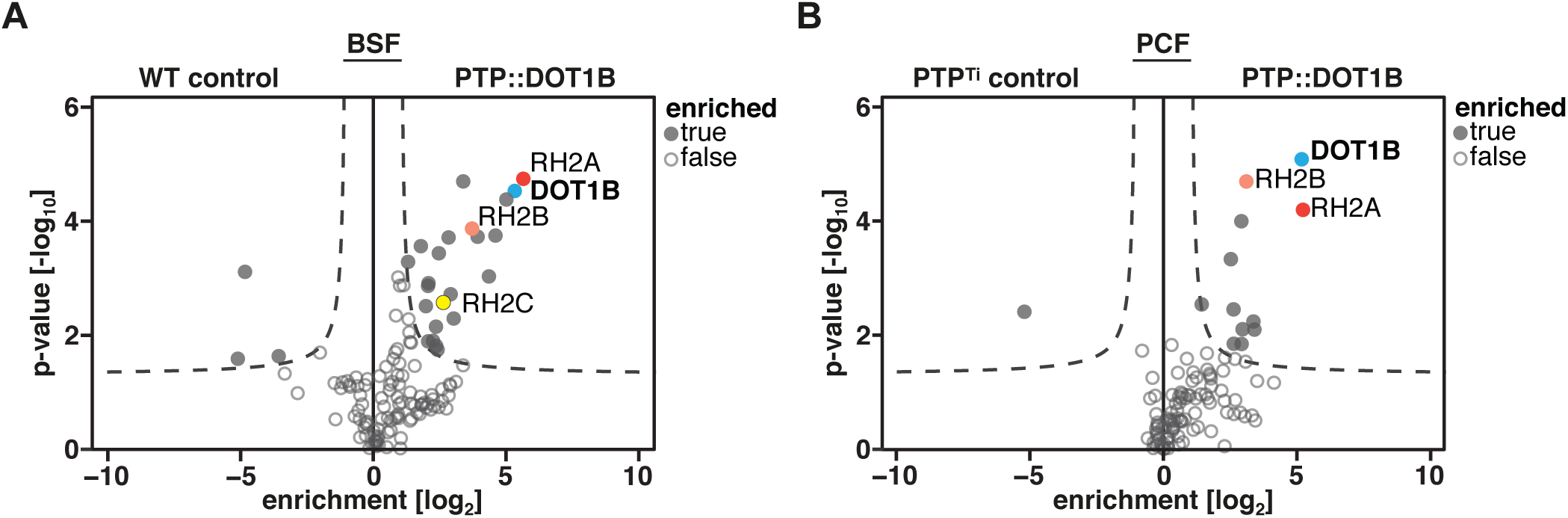
Candidate DOT1B-interacting proteins in BSF and PCF trypanosomes. Volcano plots of MS data showing proteins co-purifying with DOT1B from (**A**) mammalian-infective BSF and (**B**) the vector-specific PCF trypanosomes. The x-axis of the volcano blot represents the log2 fold-change of detected proteins in PTP::DOT1B lysates compared to control cell lysates and the y-axis shows the p-value (Welch t-test) of the four biological replicates. One of the most prominent novel interactors identified in both life cycle stages was the RH2 complex (subunits RH2A, RH2B, RH2C). Complete datasets are in Supplementary Tables S1 and S2.

Unexpectedly, we co-purified all three subunits of the RH2 complex (RH2A, RH2B and RH2C) (75), very recently described in trypanosomes by Briggs et al. (13). There are two types of RH in eukaryotes (RH1 and RH2) with overlapping functions in genome stability mediated by removing of R-loops (76). In contrast to RH1, RH2 is also involved in recognizing and cleaving single ribonucleotides falsely incorporated in DNA in a process called ribonucleotide excision repair (RER) (77). Both trypanosomal RH enzymes participate in antigenic variation by processing R-loops (12,13). Furthermore, RH2 has a role in RNA Pol II transcription in *T. brucei* (13).

To determine whether these interactions are specific for the mammalian-infective stage, we also carried out the affinity purification of DOT1B in the vector-specific procyclic form (PCF), and analyzed the results by mass spectrometry (MS). In this instance, we used an improved control cell line, which expressed the PTP-tag alone (Supplementary Figure S2B). Eleven proteins were significantly enriched together with DOT1B in the PCF PTP::DOT1B TAP, and one was significantly enriched in the control TAP (Figure 1B, Supplementary Table S2). Remarkably, we again purified two members of the RH2 complex.

In summary, we identified for the first time candidate DOT1B-interacting proteins. Moreover, the results indicate that the association of the RH2 complex with DOT1B is not stage-specific. Interestingly, all of the DOT1B co-purified nuclear proteins are potential novel interactors, as to our knowledge there are no homologs to these candidates in the list of DOT1 interactors from other organisms. We decided to focus further analysis on the most prominent candidate, the RH2 complex.

### Verification of the DOT1B-RH2 interaction

In order to validate the interaction between DOT1B and RH2, we performed reciprocal co-immunoprecipitations (co-IP) with HA-tagged RH2 subunits. Both alleles of *RH2A* were endogenously tagged at the 3’ end using homologous recombination (Supplementary Figure S3A). Integration of the targeting constructs at the endogenous loci was confirmed by PCR analysis of genomic DNA (Supplementary Figure S3B). Cells expressing the two tagged alleles grew at a comparable rate to controls, demonstrating that the *RH2A::HA* alleles were functional (Supplementary Figure S3C). The cell line expressing RH2A-HA and control WT cells were used in a co-IP experiment with pre-immobilized HA-antibody sepharose. The successful enrichment of RH2A-HA was monitored by WB analysis using anti-HA antibodies (Supplementary Figure S3D). MS analysis of eluates identified 17 proteins significantly enriched with RH2A, including both other RH2 subunits as well as DOT1B, confirming the interaction of DOT1B with the RH2 complex (Figure 2A, Supplementary Table S3). The same IP-MS approach was applied to the other RH2 subunits. While endogenous HA-tagging of the *RH2B* subunit alleles could be confirmed by diagnostic PCR, we could neither detect nor purify this fusion protein biochemically (data not shown). Tagging of the *RH2C* alleles was similarly confirmed by diagnostic PCR (Supplementary Figure S4A, B). In this instance however, the RH2C-HA protein could be readily immunoprecipitated (Supplementary Figure S4C). Mass spectrometry analysis of co-immunoprecipitating proteins yielded an enrichment of 77 candidates, including the other subunits H2A and H2B and again DOT1B (Figure 2B, Supplementary Table S4). Apart from the RH2 subunits and DOT1B, the only other overlapping nuclear candidate enriched in both co-IPs was DOT1B’s paralog DOT1A (56).

**Figure 2.**
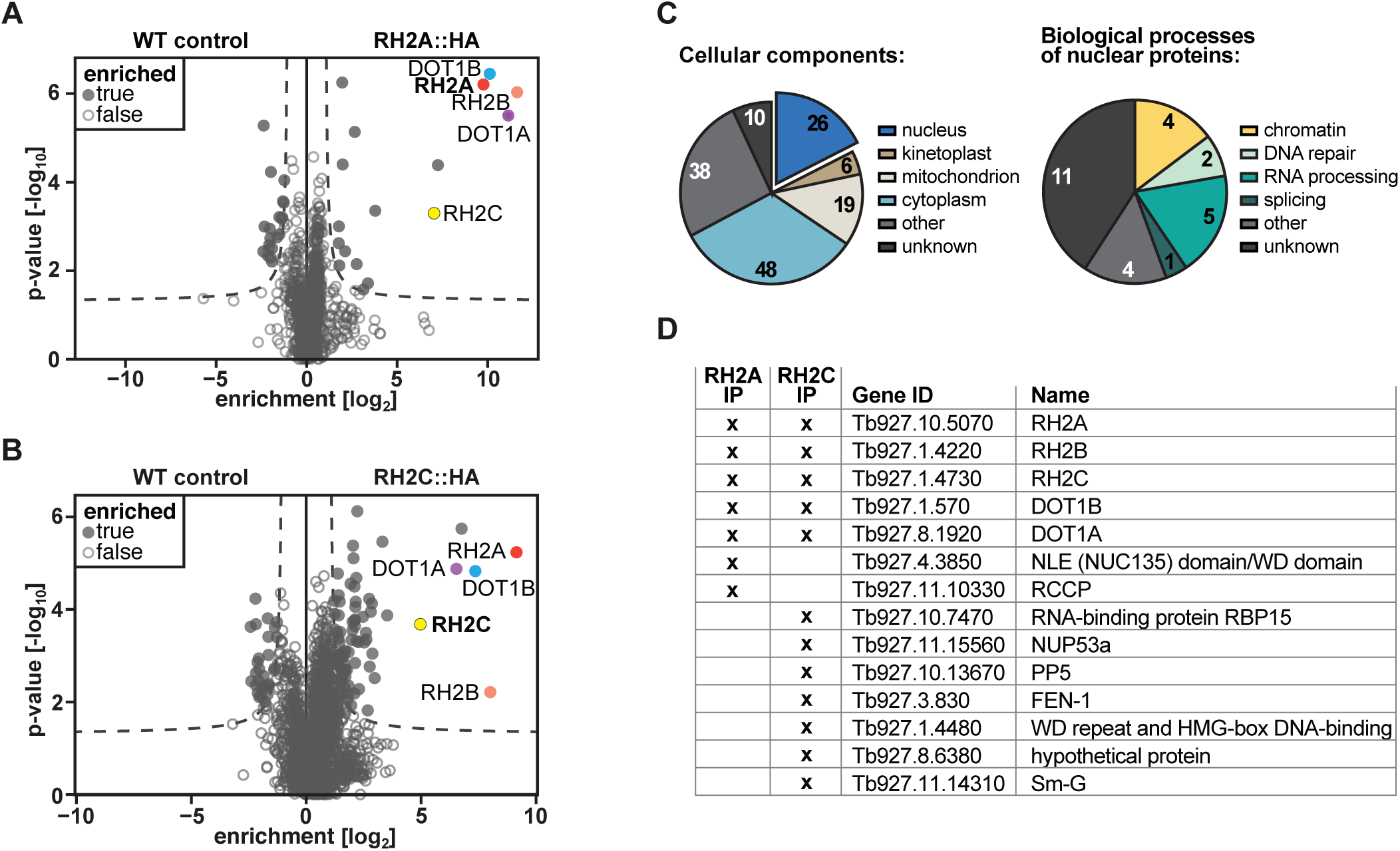
Candidate RH2-interacting proteins in PCF trypanosomes. Volcano plot of co-purified proteins after (**A**) RH2A-HA vs. WT control or (**B**) RH2C-HA vs. WT control IPs, obtained by the MS analysis of four biological replicates each. In the case of RH2A-HA, the 17 significantly enriched proteins included all subunits of the RH2 complex as well as DOT1B and DOT1A. The 77 significantly-enriched proteins of the RH2C-HA fractions included the bait protein RH2C with the other subunits RH2A and RH2B, as well as DOT1B and DOT1A. (**C**) Pie charts with the numbers of enriched proteins localizing to the listed cellular components (GOCC), as well as the numbers of enriched core proteins in the listed biological processes (GOBP). Proteins can be assigned to more than one cellular component/biological process. (**D**) Shortlist of nuclear RH2 co-enriched proteins.

We combined the candidates obtained from the subunit A and subunit C purifications and categorized them according to their location based on curated GO terms (TriTrypDB) (Figure 2C). Since it has been shown that RH2A is a core protein of a nuclear complex (13), we focused on the 26 nuclear proteins annotated (Supplementary Table S5). We further categorized the nuclear proteins into the different predicted or curated biological processes (Figure 2C, Supplementary Table S5). These proteins are mainly involved in RNA processing and DNA repair but there were, besides DOT1B, other chromatin-associated proteins such as DOT1A (56) or the ISWI-complex protein RCCP (24). Additionally, a potential Flap endonuclease 1 (FEN-1) was enriched. FEN-1 is involved in several mechanisms of RNA processing like RER, Okazaki fragment processing, or stalled replication fork rescue in other eukaryotes (78). In *T. cruzi*, FEN-1 participates in DNA replication and repair (79). FEN-1 was also described in an iPOND (isolation of proteins on nascent DNA) MS screen to be associated with replication in *T. brucei* (80). Furthermore, the spliceosome component Sm-G was enriched with RH2 (81). A shortlist of these more promising candidates was compiled (Figure 2D).

### DOT1B-mediated proximity labeling identifies 152 near neighbors

To further validate potential DOT1B-interacting proteins identified by TAP and screen for candidate DOT1B binding partners under native conditions, we next carried out proximity-dependent biotin labeling (BioID). This technique is based on the fusion of a protein of interest to the modified bacterial BirA* biotin ligase and has the advantage that the bait protein can function under native conditions throughout the cell cycle, while biotinylating all nearby proteins in a proximity-dependent fashion. Biotinylated proteins can then be purified with high stringency using streptavidin beads (82). We induced the expression of a DOT1B-Myc-BirA* fusion protein for 24 h and incubated PCF cells with 50 µM biotin for another 24 h. Microscopy analysis of DOT1B-Myc-BirA*-expressing cells using fluorescent avidin revealed a strong biotinylation signal in the nucleus, which was absent from controls (Supplementary Figure S5A). Biotinylated proteins and associated interaction partners were purified in quadruplicates using streptavidin-coated beads, and uninduced cells served as a control. WB analysis of eluate fractions of DOT1B-Myc-BirA*-expressing cells using streptavidin revealed additional bands compared to the uninduced control (Supplementary Figure S5B). Eluted fractions were also analyzed by MS. In total, we identified 152 potential near neighbors, including eleven directly-biotinylated proteins (Figure 3A, Supplementary Table S6). All RH2 subunits were significantly enriched, notably by direct biotinylation of subunit A. 79% of the identified proteins were classified as nuclear (based on their curated GO components published on TriTrypDB), including proteins predicted or known to be involved in replication, RNA processing, and transcription (Figure 3B).

**Figure 3.**
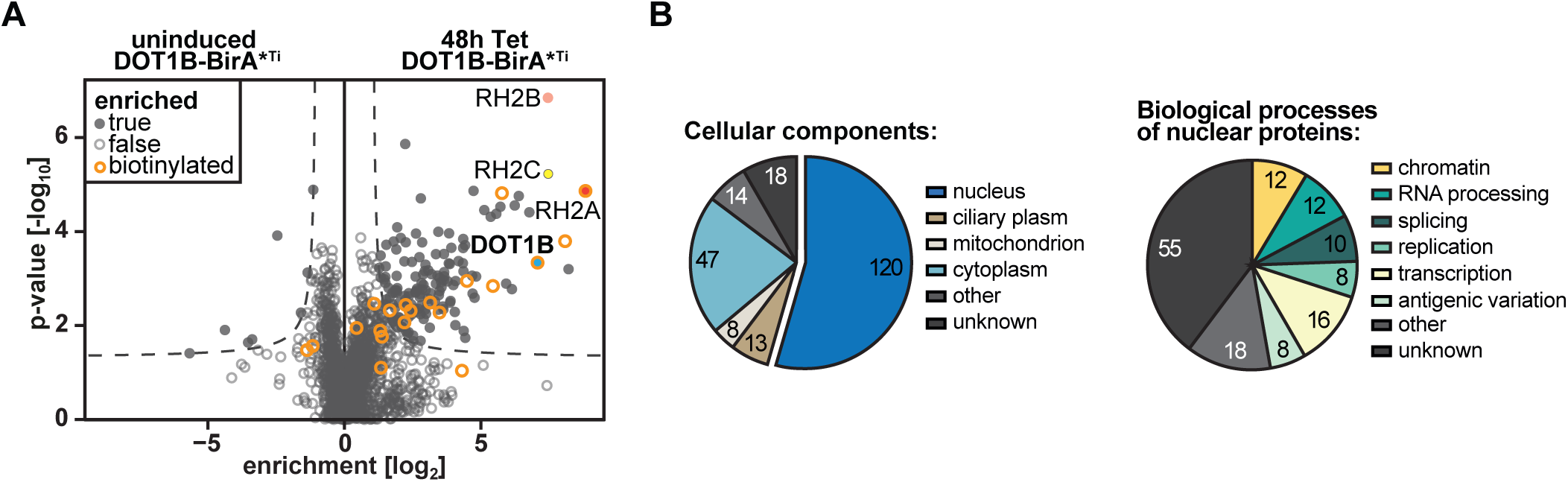
Proximity labeling with DOT1B. (**A**) MS analysis of proteins detected after streptavidin pulldowns from DOT1B-BirA*-expressing cells in the presence of excess biotin. Pulldowns from uninduced DOT1B-BirA* cells served as a control. Experiments were conducted in four biological replicates. 152 significantly-enriched proteins were identified, including all subunits of the RH2 complex. The RH2 subunit A was one of eleven directly-biotinylated proteins detected (orange circled). (**B**) Pie charts showing the number of enriched proteins in the respective cellular components (GOCC), as well as the number of enriched core proteins in the listed biological processes (GOBP). Proteins can be assigned to more than one cellular component/biological process.

Even though we could not confirm any other proteins previously-identified by TAP to be DOT1B-associated than the RH2 subunits, we have expanded the list of potential DOT1B interaction partners with other candidates. In addition, we have confirmed an association between DOT1B and the RH2 complex under native conditions, because all RH2 subunits could be identified. Interestingly, we could only detect biotinylation of the A subunit, which suggests that this subunit was closest to DOT1B and provides the interface between DOT1B and the RH2 complex. Yeast 2-hybrid data supported the assumption that subunit A of RH2 interacts directly with DOT1B (Supplementary Figure S6).

These results, along with all the other experiments mentioned above, support a direct interaction between DOT1B and the RH2 complex, most likely with the RH2A subunit forming the interface of this novel complex.

### Contribution of DOT1B to R-loop resolution

We next wanted to understand the function of the DOT1B-RH2 interaction. Depletion of subunit A of the RH2 complex leads to cell cycle stalling with accumulation of R-loops and DNA damage (13). Both phenomena were mapped to transcription initiation sites of RNA pol II, as well as to the active and inactive ES. As a consequence, the expression of several genes (including *VSG*s) was altered, which led to changes in the surface coat composition. Since *VSG*s throughout the repertoire were derepressed, recombination-based switching events were suggested. In addition, monoallelic expression was impaired, which resulted in the simultaneous expression of two VSGs on the surface of the mutant parasites (13).

In contrast, DOT1B is not essential for parasitic viability (56), indicating that the RH2 protein complex very likely also possesses DOT1B-independent functions. Although transcription-associated proteins were enriched in the DOT1B BioID, no previous experimental data has suggested an influence of DOT1B on RNA Pol II expression to date. Derepression of silent ES-associated VSGs and slow ES switching kinetics have been observed in DOT1B-depleted cells, however (27). Alterations in recombination-based switching have not been investigated. An overlapping contribution of both proteins to antigenic variation could be via regulation of genome integrity. We therefore asked whether DOT1B might support RH2A in minimizing DNA damage caused by R-loop accumulation, which might be a novel route to initiate recombination-based VSG switching (12,13,15). To test this hypothesis, we analyzed R-loop formation, DNA damage accumulation, and VSG switching in DOT1B-depleted cells.

First, we compared genome-wide R-loop formation in DOT1B-depleted cells with WT cells in a quantitative dot blot analysis (Figure 4A). Genomic DNA was analyzed using the RNA-DNA hybrid-specific S9.6 antibody (83). DOT1B RNAi cells were analyzed instead of DOT1B KO cells in order to capture early events after protein depletion. DOT1B depletion upon induction of RNAi was confirmed by monitoring loss of DOT1B-specific H3K76me3 methylation and the concomitant increase of H3K76me2 methylation by WB (Supplementary Figure S7A). To have a suitable endogenous positive control for this assay, we included DNA from RH2A-depleted cells. RNAi against RH2A resulted in a growth defect (Supplementary Figure S7B) as previously described (13). As an additional positive control, synthetic RNA-DNA hybrids were generated by first strand synthesis reaction using reverse transcriptase and *T. brucei* whole-cell RNA extracts as template. S9.6 antibody specificity was also verified by digestion of half of the samples with recombinant *E. coli* Ribonuclease H (*Ec*RH), which specifically cleaves RNA in RNA-DNA hybrids (84). Strikingly the quantification of the signals clearly demonstrated a 2.5-fold enrichment of R-loop structures in 72 h DOT1B-depleted cells compared to WT cells, suggesting that DOT1B is involved in R-loop processing (Figure 4B). The signal enrichment was observed in both 48 h RH2A-depleted as well as in cDNA samples. *Ec*RH sensitivity confirmed the specificity of the assay (Figure 4A, lower blot).

**Figure 4.**
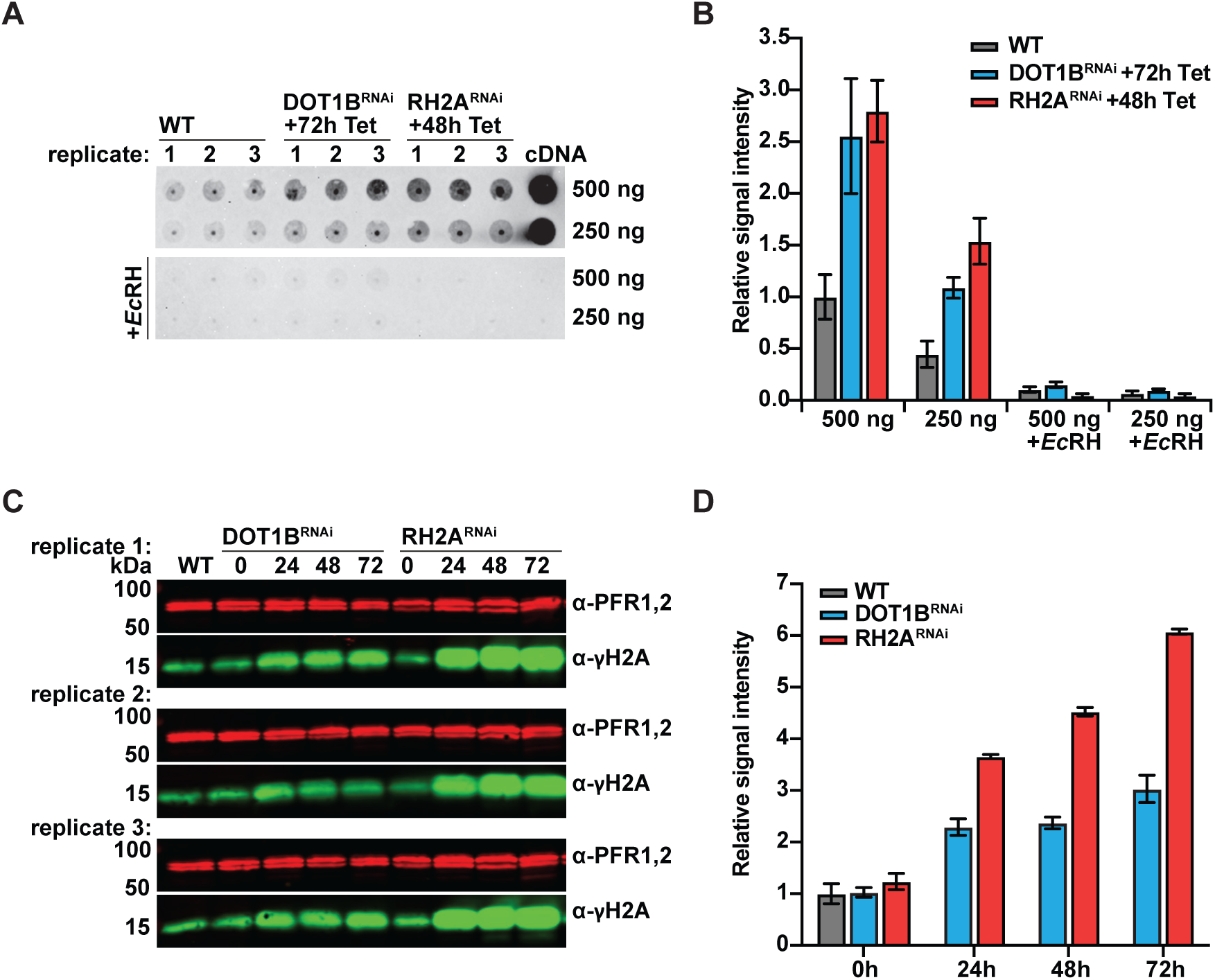
Accumulation of R-loops and DNA damage after DOT1B depletion. (**A**) R-loop dot blot of DOT1B- and RH2A-depleted cells in comparison to the WT. Protein depletion by RNAi was induced in case of DOT1B for 72h and in case of RH2A for 48h by addition of tetracycline (Tet). Genomic DNA was isolated from biological triplicates of each cell line and 500 ng and its 1:1 dilution were spotted onto a positively-charged nylon membrane. The membrane was probed with anti-RNA-DNA hybrid S9.6 antibodies. Treatment of samples with recombinant *Ec*RH was used as negative control. Detection of synthetic RNA-DNA hybrids, generated by copy DNA (cDNA) synthesis, served as positive control. (**B**) Quantification of dot blot data. WT level was set to 1. Error bars represent the standard deviation of mean values of the three biological replicates. (**C**) Western blot of the DNA damage marker γH2A at different timepoints after DOT1B and RH2A depletion by RNAi. (**D**) Quantification of the WB data. γH2A levels were normalized to PFR1,2 protein expression. WT level was set to 1. Error bars represent the standard deviation of mean values of the three biological replicates.

We next wanted to examine whether there is also an accumulation of DNA damage in DOT1B-depleted cells. To test this, we monitored DNA damage after RNAi-mediated depletion of DOT1B in quantitative WB analysis using an antibody specific for phosphorylated histone H2A (γH2A), which is a marker for DNA damage (Figure 4C) (85). Whole cell lysates from cultures 0 h, 24 h, 48 h and 72 h post-induction of DOT1B RNAi were compared to whole cell lysates of WT cells. An increased γH2A signal has previously been observed in RH2A-deficient cells (13), so whole cell lysates 0 h, 24 h, 48 h and 72 h after RH2A RNAi induction were used as positive controls. The γH2A signal was normalized to paraflagellar rod (PFR1,2) protein levels. The quantification revealed an accumulation of DNA damage in DOT1B-depleted cells (Figure 4D). This damage accumulated over time and resulted in a 3-fold increase after 72 h of RNAi induction in comparison to uninduced and WT samples. Interestingly, the accumulation of damaged chromatin appears to be less dramatic compared to the 6-fold increase of DNA damage after 72 h of RH2A subunit depletion.

We had previously observed massive accumulation of DNA damage in DOT1B-deficient cells during differentiation from BSF into PCF trypanosomes, but not in DOT1B KO BSF cell culture (57). The assumption was that damage accumulation in DOT1B mutants was differentiation-dependent. To assess whether these observations were based on overlooked compensation/adaptation processes after prolonged culture of DOT1B KO cells, we reanalyzed DNA damage in a freshly-generated KO cell line. A new KO cell line was generated in BSF trypanosomes by replacement of both alleles with drug selection markers in two consecutive transfections. Amplification of the *DOT1B* locus by PCR and analysis of H3K76me3 and H3K76me2 signals in WB confirmed the deletion of DOT1B (Supplementary Figure S8A, B). The new DOT1B KO cells were cultured for a period of 36 days post-transfection of the cassette that replaced the second allele of *DOT1B*. Whole cell lysates of WT and DOT1B KO cells after 14 days, 21 days, 28 days, and 35 days post-transfection were prepared and analyzed in a quantitative γH2A WB. PFR1,2-normalized γH2A signals were 2.8-fold increased at the 14 days timepoint in the DOT1B KO samples compared to WT samples. The γH2A signal then declined in subsequent timepoints, from the initial 2.8-fold increase after 14 days to 1.7-fold after 35 days (Supplementary Figure S8C). This supports the hypothesis that trypanosomes can partially compensate the loss of DOT1B by an unknown adaptation process.

The initial increase of DNA damage in DOT1B KO cells (Supplementary Figure S8C), and the comparable amount (3-fold) of DNA damage after depletion of DOT1B by RNAi (Figure 4C), supports the hypothesis that DOT1B influences genome integrity. Individual DSBs in subtelomeric regions of the active or inactive ES, as well as at chromosome-internal loci, are differentially tolerated by trypanosomes (8,17,71,86-88). Cells show divergent growth outcomes depending on the life cycle stage as well as the extent and location of the damage, which can be explained in part by different cell cycle responses or survival rates. Given that DOT1B is important for genome integrity, we asked whether DOT1B-deficient cells also show growth perturbations. We compared the growth of DOT1B RNAi cells after tetracycline induction to uninduced and WT cells. DOT1B-depleted cells showed a slight delay as the depletion time progresses (Supplementary Figure S7C). In contrast, DOT1B null mutants showed a severe growth defect after KO generation that gradually reverted back to WT rates over a five-week culturing period, similar to the amount of DNA damage (Supplementary Figure S8D).

### DOT1B depletion causes VSG deregulation

Since DOT1B loss or depletion, like RH2A depletion, affects R-loop levels and is accompanied with reduced genome integrity, we wanted to investigate DOT1B’s contribution to the regulation of VSG switching. First, the expression of cell-surface VSGs was analyzed in DOT1B RNAi cells by flow cytometry with a monoclonal antibody specific for the predominant VSG-2 isoform. As a control, VSG-13-expressing PN13 cells were included (27). WT cells and uninduced DOT1B RNAi cells were VSG-2 positive, whereas the PN13 cells were VSG-2 negative (Figure 5A). After 72h of DOT1B depletion, 7% of the parasites no longer expressed the original VSG-2, suggesting that these cells switched on another *VSG* gene (Figure 5A, fourth panel).

**Figure 5.**
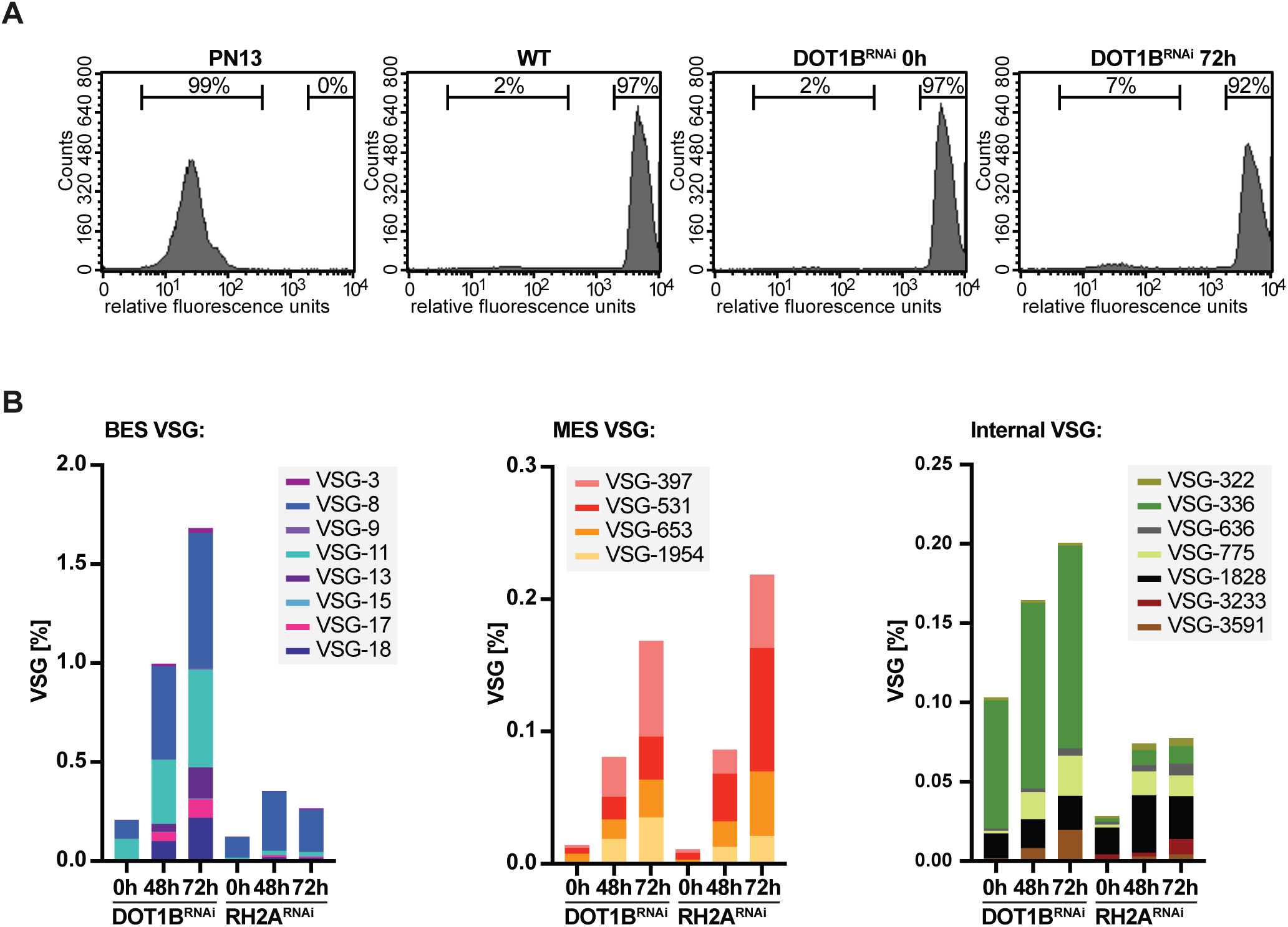
Increased expression of multiple VSGs after DOT1B depletion. (**A**) Quantitative analysis of VSG-2 expression by flow cytometry after DOT1B depletion. Uninduced cells as well as the parental cell line served as positive controls for VSG-2 expression, and the VSG-13 expressing PN13 cell line served as a negative control. 7% of DOT1B cells were VSG-2 negative after 72 h of RNAi induction. Representative profiles of one exemplary replicate per cell line are shown, percentages represent the mean values of three replicates. (**B**) Enrichment of multiple silent VSGs in induced DOT1B and RH2A RNAi cells compared to the uninduced controls. VSGs were isolated from the cell surface of four biological replicates each and analyzed by MS. The abundance of deregulated VSGs (excluding VSG-2) is given as an average percentage of LFQ intensity for each condition. BES-associated VSGs, MES-associated VSGs, as well as VSGs from internal genome loci are displayed in separate diagrams with adapted Y-axis.

To examine which *VSG*s are expressed in the VSG-2 negative parasite population, we released soluble VSGs from the cell surface of WT cells and DOT1B RNAi cells 0 h, 48 h, and 72 h after RNAi induction and analyzed them by MS. The protein level of 17 VSGs was more than 2-fold upregulated compared to WT cells after 72 h of DOT1B RNAi induction (Figure 5B, Supplementary Figure S9). Almost half of those VSGs were associated with previously-silent BSF ESs (4). In addition, four were detected which are normally only expressed from metacyclic ESs (MES) (89). Strikingly, we also detected several VSGs that are usually located in the subtelomeric arrays of the megabase chromosomes, such as VSG-636, VSG-775 or VSG-3591. We further analyzed the VSG expression pattern after different culture timepoints of the newly-generated DOT1B KO cells relative to WT cells by MS analysis of whole cell lysates. Almost the same set of deregulated VSGs was observed in early timepoints after DOT1B deletion as in the DOT1B-depleted RNAi cells (Supplementary Figure S8E, Supplementary Table S7). At late timepoints after recovery from the DNA damage phenotype, only ES-associated VSG could be detected.

In addition, we compared DOT1B RNAi with RH2A RNAi cells to assess whether the proportions of the expressed VSG repertoire are similar, further suggesting a shared regulatory function of both proteins in antigenic variation. The abundance of 15 surface VSGs was 2-fold upregulated 72 h after depletion of RH2A compared to the expressed VSGs of WT cells. Similar to DOT1B-depleted cells, VSGs associated with BES and MES, but also VSGs from subtelomeric regions were detected after RH2A depletion (Supplementary Figure S9, Supplementary Table S8). These VSGs could only have been expressed after recombination into an active BES. Strikingly, in both cell lines 80% of the expressed VSGs were identical 72 h post induction of RNAi, although the percentage of new VSGs in DOT1B-depleted parasites was higher compared to RH2A.

Taken together, our data revealed a novel interaction between DOT1B and RH2 that exists in the mammalian-infective BSF and in the insect PCF of trypanosomes. Since DOT1B-deficient cells showed increased R-loop formation, compromised genome integrity, and a similar deregulated VSG pattern as observed after RH2A depletion, we propose a common function for this complex in regulating antigenic variation by modulation of R-loop structures.

## DISCUSSION

DOT1 and the histone modifications it mediates are highly conserved across many species. In yeast and humans, knowledge about DOT1 regulation as well as its contribution to transcription, meiotic checkpoint control, and the DNA DSB response was increased substantially by the discovery of DOT1-binding factors. In contrast, nothing is known about DOT1-associated proteins in *T. brucei*. DOT1B and other factors involved in chromatin modifications, transcriptional regulation, or telomere structure maintenance were shown to influence antigenic variation in trypanosomes, but our understanding of the compositions and interplay of these molecular machineries remains incomplete. The identification of a DOT1B interactome would improve our understanding of the tightly regulated process of antigenic variation.

In this study, we identified DOT1B-associated proteins using a combination of complementary methods in *T. brucei*. TAP detected 23 proteins to be significantly enriched in BSF and 12 in PCF, whereas 152 proteins were enriched due to proximity labeling with DOT1B in PCF. The list contains proteins involved in RNA processing, including splicing, as well as replication and transcription. One of the most abundant interactors and the only overlap between all individual data sets was the RH2 complex with its three subunits. The minor overlap of the two complementary approaches may indicate many weak or transient interactions of candidates making them prone to dissociate during the affinity purification step. Alternatively, the substantially larger number of candidates detected in the BioID screen might reflect an ubiquitous presence of DOT1B at chromatin, since DOT1B is likely to be responsible for the entire trimethylation of H3 during the cell cycle and therefore probably in close proximity to many other chromatin-related proteins, such as those involved in replication and transcription. Although we tried to rule out non-specific binding of proteins to the PTP-tag by ectopic expression of the tag in our control cell line, often-reported contaminants such as ribosomal subunits were present in our enriched fractions. Hence, further experiments are needed to validate the list of remaining candidates. Reciprocal IPs and yeast 2-hybrid analysis were both used to validate and to define the mode of interaction between DOT1B and trimeric RH2 subunits in this study.

Our data did not include previously-described DOT1 nuclear interactors found in other species, such as members of the ENL family that are among the best-studied interactors of mammalian DOT1 (33,34,38,90,91). Searching the *T. brucei* proteome with human ENL or AF9 by BLAST revealed a potential YEATS protein homolog, but this protein was not significantly enriched in our assays, suggesting that this interaction is likely not conserved in trypanosomes. The conserved YEATS domain is important for targeting ENL/AF9 to chromatin through its binding to acetylated lysines of nucleosomes with high affinities to H3K9ac and H3K27ac (92-94). These interactions influence the efficiency of DOT1L to mediate di- and trimethylation marks by increasing the residency time at specific sites to produce higher methylation states (34). However, this specific regulatory process seems unlikely in trypanosomes, because the generation of di- and trimethylation of H3K76 in trypanosomes is ensured by the division of this task between two enzymes, DOT1A and DOT1B (55,56). Furthermore, a long C-terminal extension outside of the conserved HMT domain of DOT1L is essential for the interaction to DNA and other proteins, including the interaction to ENL/AF9 (33,34). Trypanosomal DOT1B only comprises the conserved HMT domain and lacks such long N- or C-terminal extensions present in yeast and human, indicating that interaction mechanisms might differ across species or might be indeed species-specific.

In this study, we identified a novel interaction between DOT1B and RH2 in BSF and PCF parasites, most likely mediated by a DOT1B/RH2 subunit A interface. To date, there are very few studies describing the mode of interactions of the RH2 complex in other organisms. In addition to a structural role to support the activity of the RH2A’s catalytic subunit, it was speculated that the other subunits might be involved in interactions with other proteins (95). Indeed, the RH2B subunit was shown to act as a platform to interact with PCNA via the conserved PCNA-interacting protein motif (so called PIP-box) (95). The only additional interactions with recruiting functions are mediated by an association of RH2A with BRCA2 (96) in humans, and telomeric proteins RIF1 and RIF2 in yeast (97). Our IPs of subunit A and C in trypanosomes revealed 26 nuclear proteins. We could not find conservation of the above mentioned interactions, but the detection of FEN1, another important player in the RER pathway downstream of RH2 (98), indicates a conserved role for RH2 in RER, a pathway that has not yet been investigated in trypanosomes. The IP of RH2B was not successful. However, it is possible that an interaction with PCNA also happens in trypanosomes. Sequence analysis in TriTrypDB suggests two putative trypanosomal RH2B isoforms and revealed that a PIP-box motif is also present at the N-terminal end of the longer RH2B isoform.

Subsequent work in this study investigated the purpose of this novel interaction with RH2 in trypanosomes. In contrast to DOT1B, RH2A is essential (13,56); hence, it is likely that RH2 has DOT1B-independent functions. As RH2 contributes to R-loop removal in trypanosomes, and increased R-loop levels are accompanied with elevated DNA damage and VSG switching (12,13), we speculated that the influence on antigenic regulation might be the overlapping function of both enzymes. Indeed, we showed that DOT1B is an additional player affecting R-loops, as R-loop formation increased in DOT1B-depleted cells. We found additional proteins described to bind to R-loops (99) and prevent their formation (100) in the BioID dataset. These include members of the spliceosome (101,102), the exosome (103), topoisomerase 1 (104) and an associated bromodomain-containing protein BRD2 (105), transcription elongation factor TFIIS (106) or FACT (107). These proteins are involved in the resolution of genome maintenance conflicts caused by R-loop formation in mammals. Interestingly, TAP of human DOT1L also identified an R-loop-interacting protein: the helicase DDX21 (108,109). Further studies are needed to evaluate whether those proteins also contribute to R-loop modulation in trypanosomes and whether this is dependent on an interplay with DOT1B.

Depletion of DOT1B results in a deregulation of VSG expression. Interestingly, we detected almost the same pattern of deregulated VSGs in DOT1B- and RH2A-deficient BSF cells. Activation of new *VSG*s does not occur randomly (110,111). *VSG* recombination events first affect BES *VSG*s, followed by subtelomeric *VSG* arrays (110). *VSG* recombination events between BES can be initiated by a break in the 70 bp repeats of the active ES and are most likely accompanied by chromatin conformational changes (6,8,71). We observed similar patterns of VSG expression deregulation as reported after break induction in the 70 bp repeats of the active ES (VSG-3, VSG-17, VSG-11) (71) or after H3.V and H4.V deletion (VSG-8, VSG-11) (6). Moreover, VSGs from subtelomeric regions (VSG-775, VSG-3591) were expressed. Both indicate that these *VSG*s may have been transferred into the active ES by homologous recombination. We propose that VSG switching may happen via inefficiently-processed R-loops, which lead to DSBs, and in turn can lead to a switching event by homologous recombination as described recently (12,13).

One possibility for how DOT1B could be involved in R-loop-driven antigenic variation could be a recruiting function of RH2A by DOT1B to R-loop sites in the genome, a task known for RH2 interactors in yeast and humans (96,97). DOT1B is responsible for fast trimethylation of nearly all newly-incorporated histones during G1-phase (our unpublished data) and therefore localizes to every domain of the entire chromatin once per cell cycle, potentially with an R-loop-processing RH2 complex as a backpack. The trimethylation of H3K76 at R-loop-free regions could generate an R-loop-processed memory in this scenario. H3K76me3 distribution assessed by ChIP in comparison to H3 occupancy and sites of R-loop accumulation (14) could be informative in this regard. Another possibility could be that DOT1B modulates RH2 activity. Interestingly, yeast RH2 was shown to perform all its functions in the G2-phase of the cell cycle (112). It would be interesting to figure out if this is also true for trypanosomal RH2, and if DOT1B contributes via a cell cycle-dependent interaction. Yeast and human subunit B and subunit C form stable sub-complexes *in vitro* (95,113), but RH2 is only active after formation of the trimeric complex with subunit A. Therefore, it could be possible that DOT1B plays a role in modulating the activity by recruiting the individual subunits to form an active complex.

Here, we provide the first evidence that DOT1B directly interacts with the RH2 protein complex. This novel physical interaction has further extended our knowledge about DOT1B’s function in antigenic variation: we have observed strikingly similar phenotypes in R-loop formation, genome integrity, and deregulation of VSGs after depletion of DOT1B or RH2A. Hence, we propose that both enzymes work together in a molecular complex to influence recombinational switching in African trypanosomes.

## ACKNOWLEDGEMENT

We thank Richard McCulloch for providing antibodies and editing the manuscript. Manfred Alsheimer for providing antibodies, Susanne Kramer for providing plasmids and Brooke Morriswood for providing plasmids and helpful comments. We thank Anja Freiwald for technical assistance and Mario Dejung for help with bioinformatics analysis. This work was supported by the CRC1361 on Regulation of DNA Repair and Genome Stability (FB) and funding from the Deutsche Forschungsgemeinschaft JA 1013/7-1 (CJJ).

