## Supplementary figures for "A DOT1B/Ribonuclease H2 protein complex is involved in R-loop processing, genomic integrity and antigenic variation in *Trypanosoma brucei*"

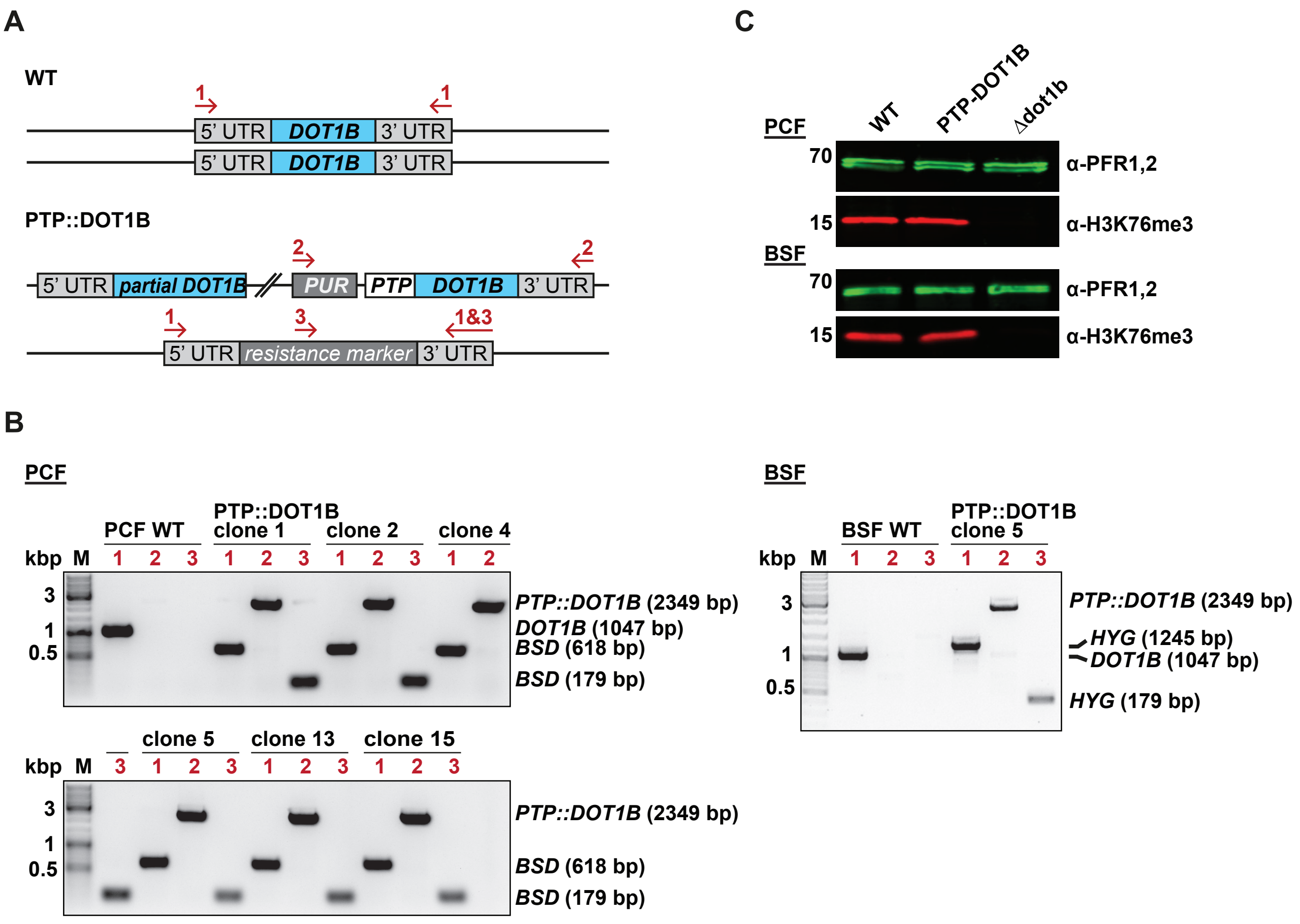


**Supplementary Figure S1.** PTP tagging of DOT1B in PCF and BSF trypanosomes. (**A**) Illustration of the endogenous *DOT1B* locus in WT and PTP::DOT1B cells. The *PTP* tag was fused to the 5' end of the first allele of *DOT1B* under puromycin (PUR) selection. The second allele of *DOT1B* was replaced by the blasticidin (*BSD*) resistance marker in PCFs, and by the hygromycin (*HYG*) resistance marker in BSF trypanosomes. Red arrows indicate the primers used for integration control by PCR. (**B**) Integration PCR with primers binding in the 5' and 3'UTR of *DOT1B* and within the resistance marker ORFs as indicated in A. Genomic DNA of six different clones were tested in PCF, one in BSF. Genomic DNA of WT cells was used as a control. Further studies in PCF were carried out with PTP::DOT1B clone 15. (**C**) Confirmation of the H3K76 trimethylation activity of the PTP-tagged DOT1B. Whole cell lysates of WT, PTP::DOT1B and Δdot1b cells were analyzed by immunoblotting with anti-H3K76me3 antibody. As a protein loading control, the same blot was probed with anti-PFR1,2 antibody.

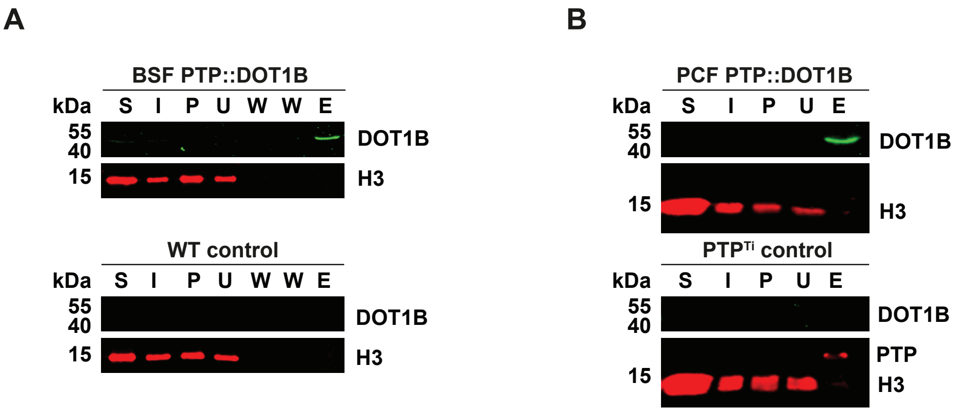


**Supplementary Figure S2.** Enrichment of DOT1B and the PTP$^{Ti}$ control after affinity purifications. Representative Western blots with samples taken during the purification procedure of (**A**) PTP::DOT1B (49.5 kDa) and the WT control in BSF or (**B**) PTP::DOT1B and PTP$^{Ti}$ control (18.6 kDa) in PCF. Whole cell lysates (S) were separated by centrifugation into soluble supernatants (I) and insoluble pellets (P). Supernatants were incubated with protein G sepharose beads. Further samples of unbound material (U), the subsequent washing steps of the beads (W), and of the proteins eluted from the beads (E) were taken. 16.5-fold more of the eluate was loaded compared to the other samples isolated during the purification procedure for the PCF pulldown and 15-fold more for the pulldown in BSF parasites. Blots were probed with anti-DOT1B antibody and anti-H3 antibody.

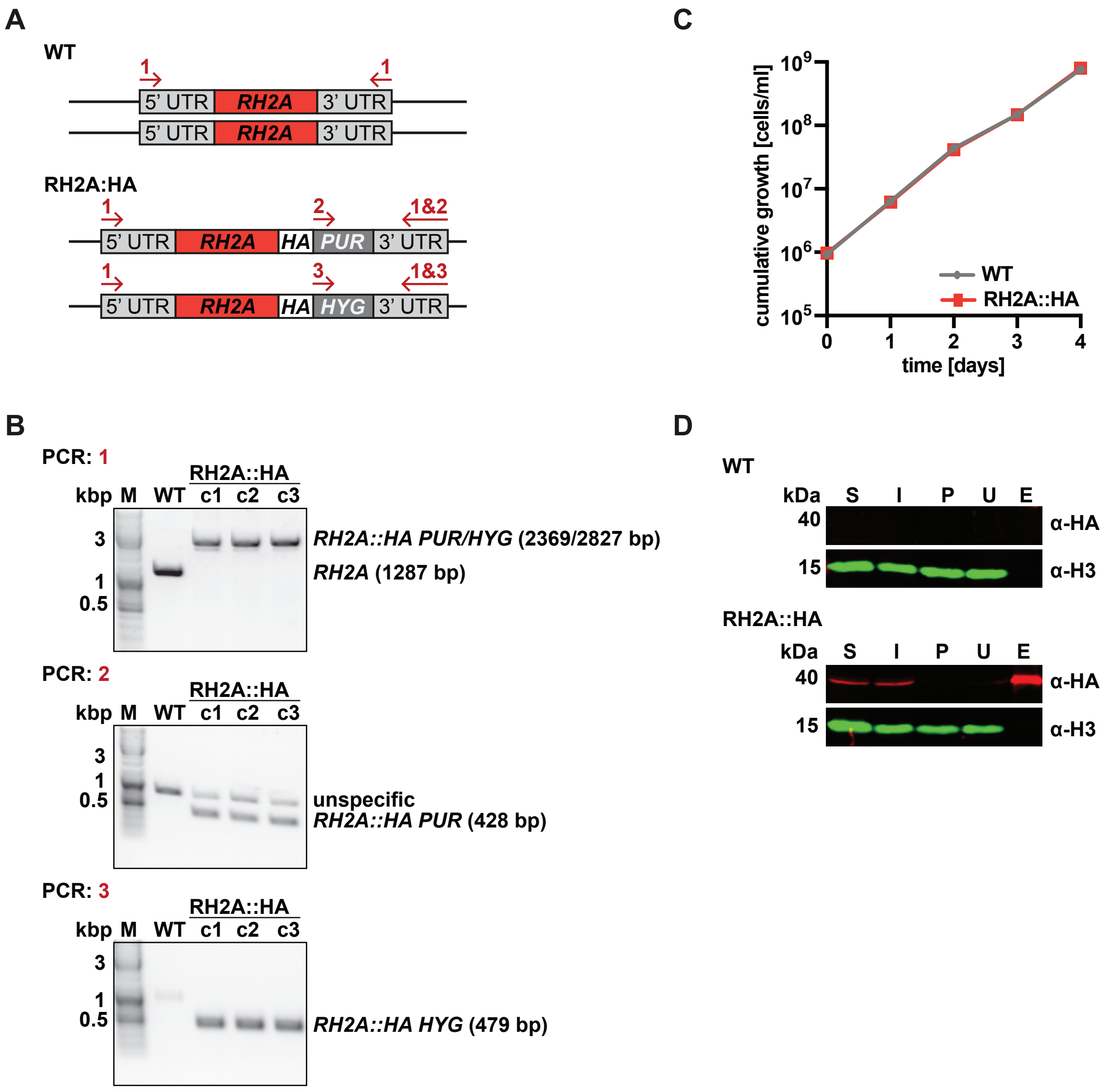


**Supplementary Figure S3.** HA tagging of *RH2A* in PCF trypanosomes. (**A**) Illustration of the *RH2A* gene locus in WT and RH2A::HA cells. The HA tag was fused to the 3' end of both alleles of *RH2A* in PCF trypanosomes. Arrows indicate the primers used for integration control by PCR. (**B**) Integration PCR with primers binding in the 5' and 3'UTR of *RH2A* and within the resistance marker ORFs, as indicated in A, verified integration of constructs. Genomic DNA of three different RH2A::HA clones was tested and genomic DNA of WT cells served as control. Further analysis in this study was carried out with RH2A::HA clone 1. (**C**) Cumulative growth shows no difference between PCF WT and RH2A::HA cells (n=3). (**D**) Representative WB of the RH2A-HA (39.3 kDa) and WT control IPs. Whole cell lysates (S) were separated by centrifugation into soluble supernatants (I) and insoluble pellets (P). Supernatants were incubated with anti-HA antibody conjugated to sepharose. Samples of unbound material (U) and of the eluates (E) were taken. 26-fold more of the eluate was loaded compared to the other samples isolated during the purification procedure. The average amount of purified RH2A-HA of the four biological replicates compared to the input material was 27.5%. The blots were probed with anti-HA and anti-H3 antibody.

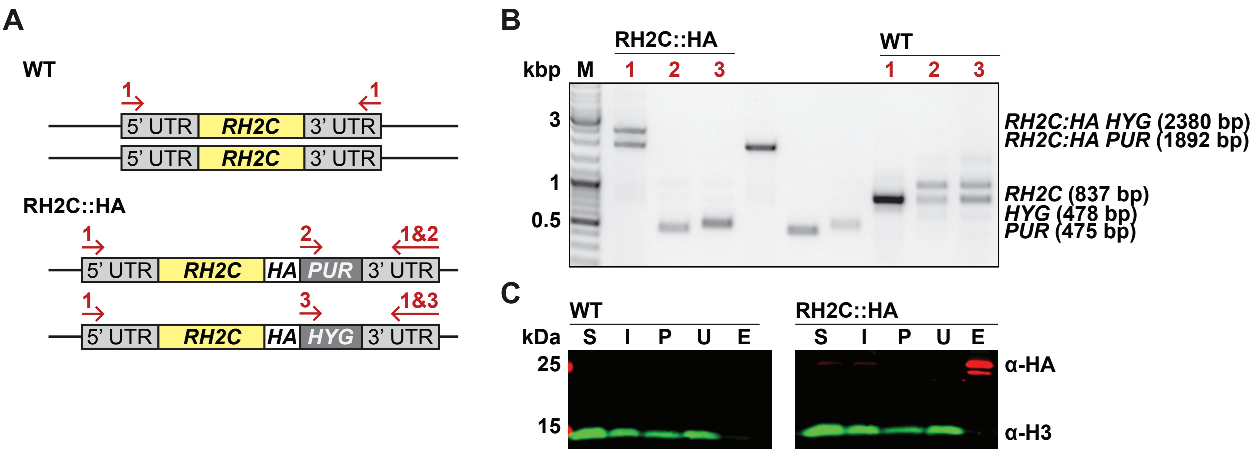


**Supplementary Figure S4.** HA tagging of *RH2C* in PCF trypanosomes. (**A**) Illustration of the *RH2C* gene locus in WT and RH2C::HA cells. The HA tag was endogenously fused to the 3' end of both alleles of *RH2C*. Arrows indicate the primers used for integration control. (**B**) Integration PCR with different primer combinations. Primers binding in the 5' and 3'UTR of *RH2C* were used to confirm the fusion of the HA tag to *RH2C*. Primers binding in the resistance marker ORFs and 3'UTR of *RH2C* confirmed 3' fusion of the tag. Genomic DNA of WT cells served as a control. (**C**) Representative WB with samples taken during the purification procedure of RH2C-HA (20.4 kDa). Whole cell lysates (S) were separated by centrifugation into soluble supernatants (I) and insoluble pellets (P). Supernatants were incubated with anti-HA antibody sepharose conjugates and samples of unbound fractions (U) and of the eluate (E) were taken. 21-fold more of the eluate was loaded compared to the other samples isolated during the purification procedure. Average RH2C-HA IP efficiency of quadruplicates was 25%. Samples were immunoblotted using anti-HA antibody and anti-H3 antibody.

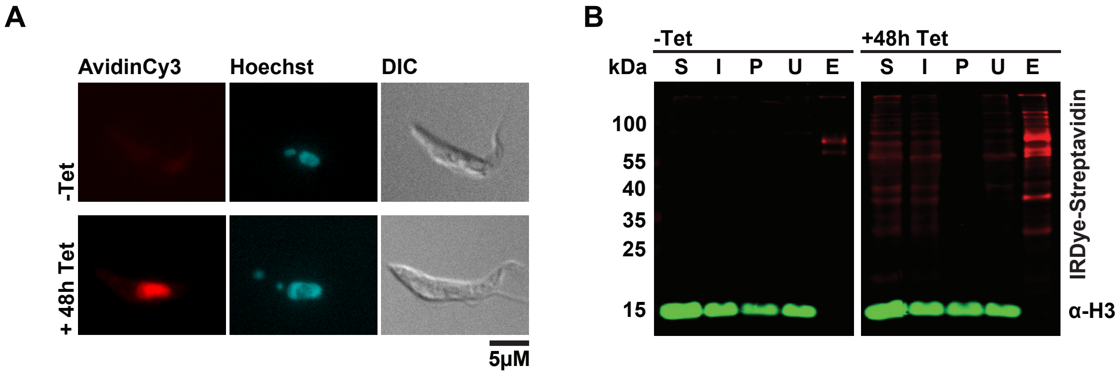


**Supplementary Figure S5.** Biotinylation of neighboring proteins by DOT1B-BirA*. (**A**) Immunofluorescence analysis of cells after incubation with biotin either with or without ectopic DOT1B-BirA* expression by addition of tetracycline (Tet). Biotinylated proteins labeled with fluorescently-conjugated avidin were observed in the nucleus. DNA was stained with Hoechst. (**B**) Representative WB with samples taken during the purification procedure after incubation with biotin of ectopically expressing DOT1B-BirA* cells or uninduced control cells. Whole cell lysates (S) were separated by centrifugation into soluble supernatants (I) and insoluble pellets (P). Supernatants were incubated with Streptavidin-conjugated agarose beads and samples of unbound fractions (U) and of the eluate (E) were taken. 38-fold more of the eluate was loaded compared to the other samples isolated during the purification procedure. Average purification efficiency of biotinylated proteins, calculated from the dominant 35 kDa protein from four replicate experiments, was 22.5%. Samples were immunoblotted using anti-H3 antibody and IRDye-Streptavidin.

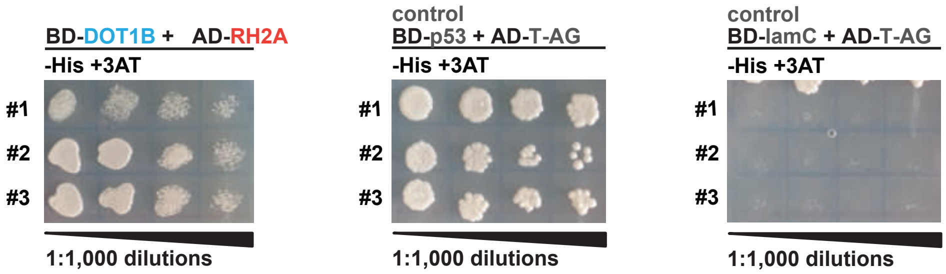


**Supplementary Figure S6.** Interaction between DOT1B and RH2A in the yeast 2-hybrid assay. Proteins of interest were either fused to the DNA-binding domain (BD) or to the activation domain (AD) of the yeast Gal4 transcription factor and were used to transform yeast cells. Growth of yeast in medium lacking histidine (-His) indicates interaction between the two proteins of interest. To reduce false positives, the stringency was adapted by growing cells in medium containing 2 mM 3-amino-1,2,4-triazole (3AT). Additionally, controls recommended by the manufacturer were probed.

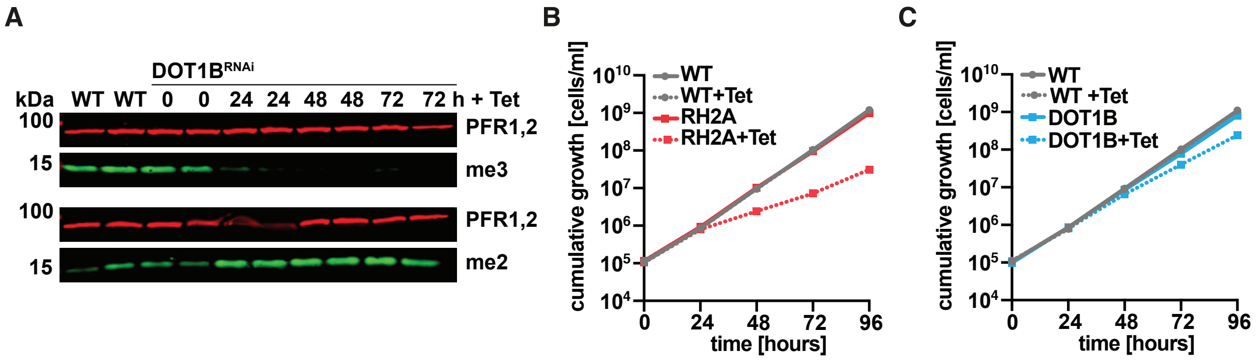


**Supplementary Figure S7.** Depletion of DOT1B and RH2A in *T. brucei* BSF. (**A**) WB with whole cell lysates taken during different timepoints after RNAi induction by addition of tetracycline (Tet), analyzed in duplicates. Decrease of DOT1B-specific H3K76me3 signal (me3) and the associated increase of H3K76me2 (me2) signal confirmed DOT1B depletion. Anti-PFR1,2 antibody was used as a loading control. (**B**) Depletion of RH2A using RNAi results in a strong growth defect. (**C**) Depletion of DOT1B using RNAi results in a mild growth defect (n=3).

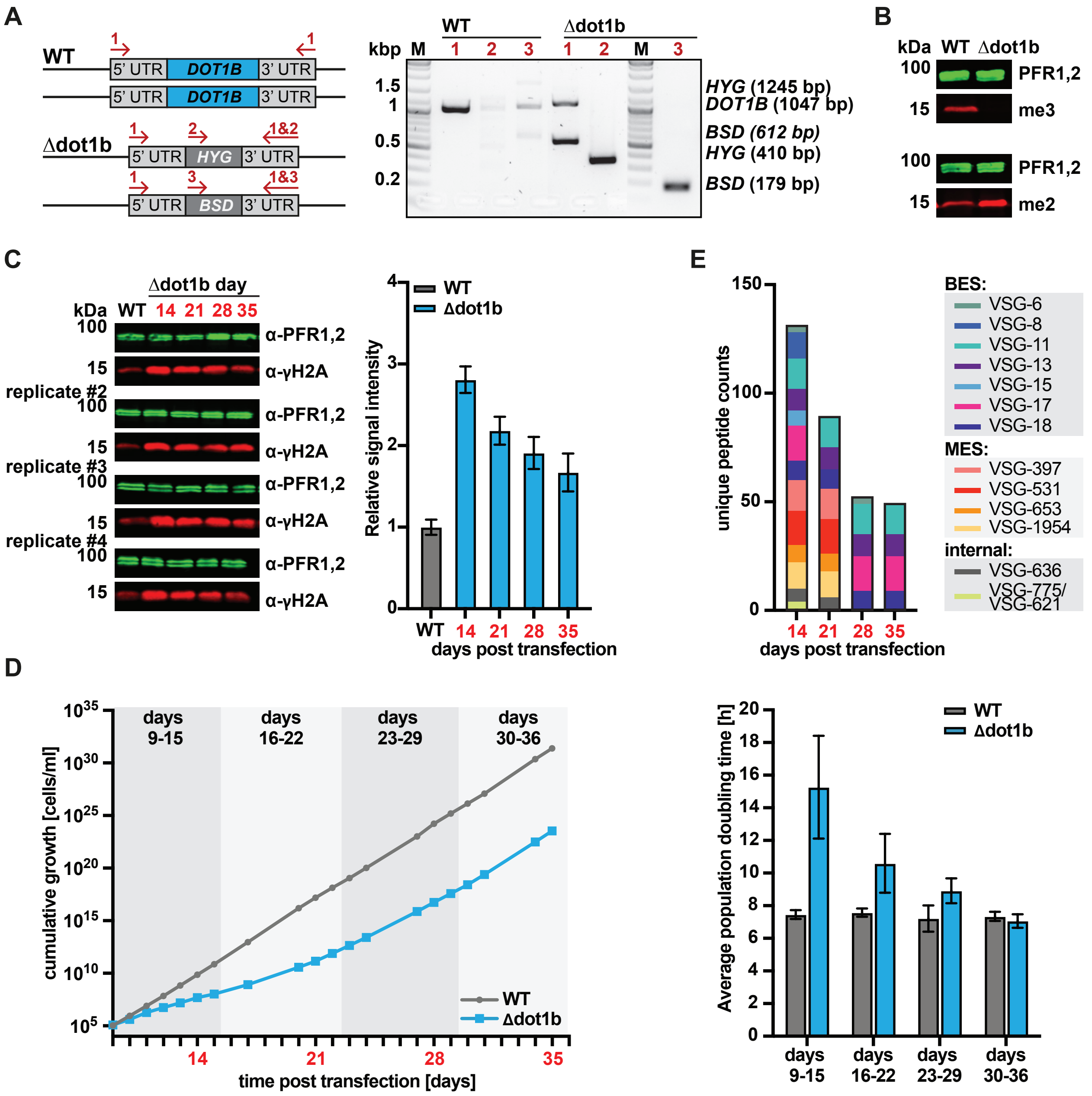


**Supplementary Figure S8.** Adaptation phenotypes of Δdot1b cells. (**A**) Illustration of the *DOT1B* gene locus in WT and Δdot1b cells. Alleles were replaced with resistance marker ORFs of hygromycin (*HYG*) and blasticidin (*BSD*). Arrows indicate the primers used for integration control. Primers binding in the 5' and 3'UTR were used to confirm the KO of *DOT1B*. Primers binding in the resistance marker ORFs and 3'UTR of *DOT1B* confirmed integration of respective markers at the right locus. Genomic DNA of WT cells served as a control M, marker lane. (**B**) Confirmation of the loss of H3K76me3 in Δdot1b cells with a corresponding increase of H3K76me2 signal. Whole cell lysates of WT and Δdot1b cells were analyzed by immunoblotting with anti-H3K76me3 and anti-H3K76me2 antibody. As a protein loading control, the same blot was probed with anti-PFR1,2 antibody. (**C**) WB and its quantitative analysis of DNA damage marker γH2A in Δdot1b cells 14, 21, 28 and 35 days post deletion of DOT1B. A reduction of DNA damage was observed over time. γH2A levels were normalized to PFR1,2 protein expression. WT level was set to 1. Error bars represent the standard deviation of the four biological replicates. (**D**) The severe growth defect of Δdot1b cells was gradually lost over a period of five weeks after generation of the KO cell line (n=4). The average population doubling time decreased over the same time period until it returned to roughly WT levels. (**E**) Mass spectrometry analysis of whole cell lysates reveals significant enrichment of multiple VSGs in Δdot1b compared to WT cells. BES-associated VSGs, MES-associated VSGs as well as VSGs from internal genome loci were deregulated during the early stages after KO generation; later only BES-associated VSGs were enriched.

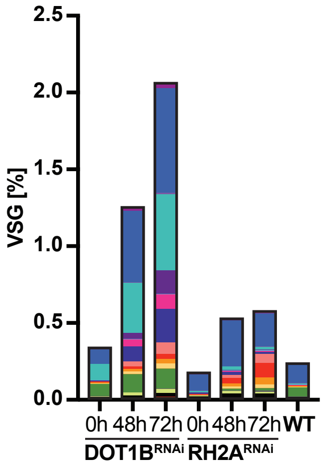


| | Localization in ES: | Localization in genome: | fold enrichment of 72h DOT1B RNAi compared to WT: | fold enrichment of 72h DOT1B RNAi compared to 0h: | fold enrichment of 72h RH2A RNAi compared to WT: | fold enrichment of 72h RH2A RNAi compared to 0h: |
|---|---|---|---|---|---|---|
| VSG-3 | BES 7 | MBC 6 | 24.2 | 15.1 | 6.2 | 3.7 |
| VSG-8 | BES 12 / BES 14 | MBC 2 / MBC 7 | 5.7 | 7.4 | 1.8 | 2.1 |
| VSG-9 | BES 2 | IC | 3.9 | 3.1 | - | - |
| VSG-11 | BES 15 | MBC 3 | 66.2 | 4.8 | 2.1 | 1.3 |
| VSG-13 | BES 17 | IC | 120.5 | 66.7 | 2.5 | 1.1 |
| VSG-15 | BES 10 | IC | 2.1 | 2.1 | 5 | 4.7 |
| VSG-17 | BES 13 | IC | 68.2 | 42 | 4.4 | 3.7 |
| VSG-18 | BES 5 | MBC 5 | 168.1 | 55.9 | 12.4 | 7 |
| VSG-397 | MES | MBC 10 | 56.9 | 38.8 | 43.5 | 31.3 |
| VSG-531 | MES | MBC 3 | 7.7 | 7.4 | 22 | 17.6 |
| VSG-653 | MES | MBC 11 | 5.6 | 4.7 | 9.7 | 35.3 |
| VSG-1954 | MES | MBC 8 | 16.3 | 20.9 | 9.8 | 12.2 |
| VSG-322 | - | UD | 2 | 1.1 | 5.1 | 3.4 |
| VSG-336 | - | MBC 8 | 2.2 | 1.6 | - | 5 |
| VSG-636 | - | MBC 9 | 4.5 | 3.1 | 6.8 | 4.2 |
| VSG-775 | - | MBC 10 | 17.4 | 14.5 | 9 | 7.3 |
| VSG-1828 | - | MBC 9 | 1.5 | 1.4 | 1.9 | 1.6 |
| VSG-3233 | - | MBC 7 | - | - | 3.1 | 3.2 |
| VSG-3591 | - | MBC 10 | 18.2 | 11.6 | 4 | 3.1 |

**Supplementary Figure S9.** Increased expression of several VSGs after DOT1B or RH2A depletion. The graph shows the percentages of VSGs identified by mass spectrometry during the different timepoints after RNAi induction of DOT1B and RH2A cell lines. The parental cell line was analyzed as a control. The table displays the genomic localization of analyzed VSGs, showing that VSGs were deregulated from throughout the genome repertoire. In addition, the fold increase of each VSG value is shown 72 hours after RNAi induction compared to the values of uninduced and WT control. Interestingly, nearly the same VSGs were deregulated in the two different cell lines. BES (BSF expression site), MES (Metacyclic expression site), MBC (megabase chromosome), IC (intermediate chromosome), UD (undefined).
