## Supplementary methods for "A DOT1B/Ribonuclease H2 protein complex is involved in R-loop processing, genomic integrity and antigenic variation in *Trypanosoma brucei*"

### SUPPLEMENTARY MATERIAL AND METHODS

#### Trypanosome cell lines

*Δdot1b*

A PCR-based gene deletion approach was used to sequentially replace both alleles of *DOT1B* with the ORFs of the resistance markers *HYG* and *BSD* in BSF SM cells. *HYG* was amplified from pHD309, *BSD* from pPOTv4 with primers containing 70 bp homology flanks of the *DOT1B* UTRs for homologous recombination. Correct replacement of the ORFs was verified by diagnostic PCR.

#### Immunofluorescence analysis

$1x10^7$ cells were fixed in 1 ml HMI-9 medium containing 2% formaldehyde (5 min, RT). Cells were washed three times with 1 ml PBS (1,000 x g, 5 min, RT) and settled on Poly-L-Lysine-coated slides. The cells were permeabilized in 0.2% Igepal CA-630/PBS (5 min, RT) and then blocked in 1% BSA/PBS (1h, RT). After removing the blocking solution, ExtrAvidinCy3 (Sigma) was applied 1:100 in 0.1% BSA/PBS including 5 µg/ml Hoechst (30 min, RT, dark). Slides were washed with PBS and mounted with Vectashield (Vecta Laboratories Inc.). Images were captured with a Leica DMI 6000B microscope and processed with the software Fiji.

#### Yeast 2-hybrid assay

The *DOT1B* and *RH2A* ORF (excluding their stop codon) were amplified from genomic DNA and cloned into the Gal4-binding domain pGBKT7 vector or Gal4-activation domain pGADT7 vector between NdeI and BamHI sites, respectively. pGADT7-T-AG and pGBKT7-p53 plasmids were used as a positive control, and pGADT7-T-AG with pGBKT7-lamC as a negative control (Clonetech).
Yeast strain AH109 was cotransformed with binary combinations of the corresponding plasmids using the lithium acetate method. Briefly, 40 ml culture ($OD_{600}$ 1.5) grown in YPDA (20 g/l Bacto Peptone, 10 g/l Yeast extract, 0.2% adenine hemisulfate, pH 5.5) were harvested (700 x g, 5 min, RT) and resuspended in 2 ml One-step-Buffer (0.2 M lithium acetate, 40% PEG3350, 100 mM DTT). Binary combinations of 1 µg bait and prey plasmids were added to 100 µl of yeast cells and incubated (30 min, 45°C). Cells were plated on tryptophan (trp) and leucine (leu) depleted SD plates to select for cotransformed plasmids (4 to 5 days, 30°C). Four to six positive clones were dissolved by rocking at 700 rpm in 100 µl PBS (15 min, RT). 10 µl of each 1:10, 1:100 and 1:1,000 serial dilution was dropped on SD selection plates (-trp, -leu) also lacking histidine (his), to select for protein-protein interactions. Stringency was increased by supplementation of SD selective plates (-trp, -leu, -his) with 2 mM 3-Aminothiazol (3AT). The plates were incubated for 3 to 4 days at 30°C. An interaction was defined as positive when growth could be observed in at least 75% of the plated clones.

### SUPPLEMENTARY TABLE LEGENDS

**Supplementary Table S1.** List of significantly enriched proteins after BSF PTP::DOT1B vs. WT control TAP obtained by the MS analysis of four biological replicates each.

**Supplementary Table S2.** List of significantly enriched proteins after PCF PTP::DOT1B vs. $PTP^{Ti}$ control TAP obtained by the MS analysis of four biological replicates each.

**Supplementary Table S3.** List of significantly enriched proteins after PCF RH2A::HA vs. WT control IP obtained by the MS analysis of four biological replicates each.

**Supplementary Table S4.** List of significantly enriched proteins after PCF RH2C::HA vs. WT control IP obtained by the MS analysis of four biological replicates each.

**Supplementary Table S5.** Combined list of significantly enriched proteins after PCF RH2C::HA IP and PCF RH2C::HA IP. Table further contains information about the cellular components (GOCC) and biological processes (GOBP) of the proteins and how they were assigned to the pie charts of figure 2.

**Supplementary Table S6.** List of significantly enriched proteins after PCF DOT1B-BirA* +Tet vs. uninduced DOT1B-BirA* control BioID obtained by the MS analysis of four biological replicates each. Table further contains information about the cellular components (GOCC) and biological processes (GOBP) of the proteins and how they were assigned to the pie charts of figure 2.

**Supplementary Table S7.** Table showing significantly deregulated proteins after different timepoints of DOT1B deletion vs. WT control obtained by the MS analysis of four biological replicates each.

**Supplementary Table S8.** List of surface VSGs at different timepoints of DOT1B and RH2 depletion by RNAi, identified by the MS analysis. WT cells (parental 2T1 cell line) were analyzed as control. Table further contains the abundance of deregulated VSGs (excluding VSG-2), given as an average percentage of LFQ intensity.
